# Structural insights into the assembly and activation of IL-27 signalling complex

**DOI:** 10.1101/2022.02.18.481027

**Authors:** Yibo Jin, Paul K. Fyfe, Scott Gardner, Stephan Wilmes, Doryen Bubeck, Ignacio Moraga

## Abstract

Interleukin 27 (IL-27) is a heterodimeric cytokine that elicits potent immuno-suppressive responses. Comprised of EBI3 and p28 subunits, IL-27 binds GP130 and IL-27Rα receptor chains to activate the JAK/STAT signalling cascade. However, how these receptors recognize IL-27 and form a complex capable of phosphorylating JAK proteins remains unclear. Here, we used cryo electron microscopy (cryoEM) to solve the structure of the IL-27 receptor recognition complex. Our data show how IL-27 serves as a bridge connecting IL-27Rα with GP130 to initiate signalling. While both receptors weakly bind the p28 component of the heterodimeric cytokine, EBI3 stabilizes the complex by binding a positively charged surface of IL-27Rα. We find that assembly of the IL-27 receptor recognition complex is distinct from both IL-12 and IL-6 cytokine families and provides a mechanistic blueprint for tuning IL-27 pleiotropic actions.

## Introduction

IL-27 is an immunosuppressive cytokine involved in resolving T cell-mediated inflammation. IL-27 inhibits Th-17 responses ^1–3^ and induces differentiation of T regulatory cells ^4^. T cell stimulation by IL-27 promotes secretion of the anti-inflammatory cytokine, IL-10, further contributing to a reduction in the inflammatory response ^5^. Together, these properties make IL-27 an attractive drug target to treat T cell-mediated inflammatory disease. IL-27 also contributes to immune exhaustion by regulating expression of co-inhibitory receptors ^6,7^. Elevated levels of IL-27 gene signatures are found in cancer and are associated with poor prognoses ^8^. Understanding how IL-27 engages its cellular receptors to form a signalling complex will provide fundamental insight into immune regulation, which could facilitate the development of new therapeutics that target IL-27 responses.

IL-27 is a heterodimeric cytokine composed of two gene products, Epstein-Barr virus-induced gene 3 (EBI3) and IL-27p28 (p28) ^9^. Grouped within the IL-12 family of heterocytokines, p28 and EBI3 share sequence homology with two components of IL-12: p35 and p40, respectively ^10^. Unlike other heterodimeric cytokines (IL-12 and IL-23) which activate Signal Transducer and Activator of Transcription (STAT) STAT3 and STAT4, IL-27 primarily induces activation of STAT1 and STAT3 pathways ^11^. These differences can be attributed to the engagement of GP130 by IL-27, which is shared across the IL-6 cytokine family ^12^.

IL-27 bridges two major cytokine families, raising the question of whether competition or synergism could occur in dimer formation between them. Under some conditions, p28 and EBI3 can be secreted independently and associate with other proteins to induce differential responses ^13–17^. Promiscuity in chain combinations is characteristic of the IL-6/IL-12 family and allows the system to produce different biologically active factors starting from relatively few precursor molecules.

GP130 binds a range of cytokines to elicit diverse cellular responses ^18,19^. The receptor has two cytokine binding sites. The first is located within the elbow between its cytokine homology regions (CHR) and has been shown to engage site 2 of cytokines, including IL-6 ^20^, Leukemia inhibitory factor (LIF) ^21^ and Ciliary neurotrophic factor (CNTF) ^22^. The N-terminal immunoglobulin (Ig) domain of GP130 comprises an additional cytokine-binding interface that recognises an epitope at site 3, resulting in higher order assemblies with different stoichiometries ^20,23^. In the hexameric GP130:IL-6 signalling complex, two molecules of GP130 dimerize to trigger JAK/STAT activation ^23^. Here, the two low affinity site 3 interfaces are necessary to stabilize the complex and trigger receptor activation by IL-6. For GP130 complexes with LIF and CNTF cytokines, site 3 is occupied by the co-receptor LIF-R ^22,24^. Structural details for how GP130 forms a heterodimeric signalling complex by binding site 3 of a cytokine remains unresolved.

Here, we report a 3.8 Å resolution cryo-EM structure of IL-27 in complex with cytokine binding domains of GP130 and its co-receptor IL-27Rα. This structure reveals a modular assembly mechanism that differs from those described for other members of the IL-12 and IL-6 families. IL-27 binds with high affinity to IL-27Rα at site 2, which is further stabilized by electrostatic interactions between IL-27Rα and EBI3. Subsequently, GP130 is recruited to the heterodimeric cytokine in a second step. Our structural and biochemical data explain how IL-27 coordinates two signalling receptors to modulate T cell mediated inflammation. Our results provide a blueprint for developing new therapeutics that target and tune IL-27 responses.

## Results

### Molecular architecture of the IL-27 cytokine recognition complex

IL-27 signals through dimerization of the cognate receptor IL-27Rα and the shared receptor GP130 ^25^. Current models for the assembly of complexes mediated by IL-27 have been largely based on structural principles derived from the IL-12/IL-23 or IL-6 systems. However, given differences in signalling outcomes between IL-27 and these two families, together with the growing importance of IL-27 as a therapeutic target, we used cryoEM to solve the structure of the IL-27 cytokine recognition complex.

To overcome challenges in cytokine stability, we engineered an IL-27 cytokine variant in which the two monomeric components (EBI3 and p28) were fused by a short linker ^25^. The fusion cytokine was expressed in insect cells and purified together with Domains 1-3 (D1, D2, D3) of GP130 and the first two domains of IL-27Rα (Supplementary Fig. 1). We then used cryoEM to solve the structure of the complete IL-27 cytokine-recognition complex (Fig. 1, Supplementary Fig. 2 and Supplementary Fig.3). We used the ab initio reconstruction protocols within cryoSPARC ^26^ to generate an initial model of the complex. Maps were further refined using a combination of homogenous and heterogenous refinement procedures. The final map was refined to a reported resolution of 3.8 Å using the gold standard FSC 0.143 cut-off, however an analysis of the angular distribution of particles indicated strong preferential orientations which impacted the quality of the map in some directions. We therefore limit our interpretation of the structure to the assignment of individual domains with models derived from Alpha fold predictions ^27^ refined with strong adaptive distance restraints and geometric restraints imposed (Supplementary Table 1). The chirality of the helical bundle central to the p28 cytokine was used to assign the handedness of the reconstruction.

**Figure 1.**
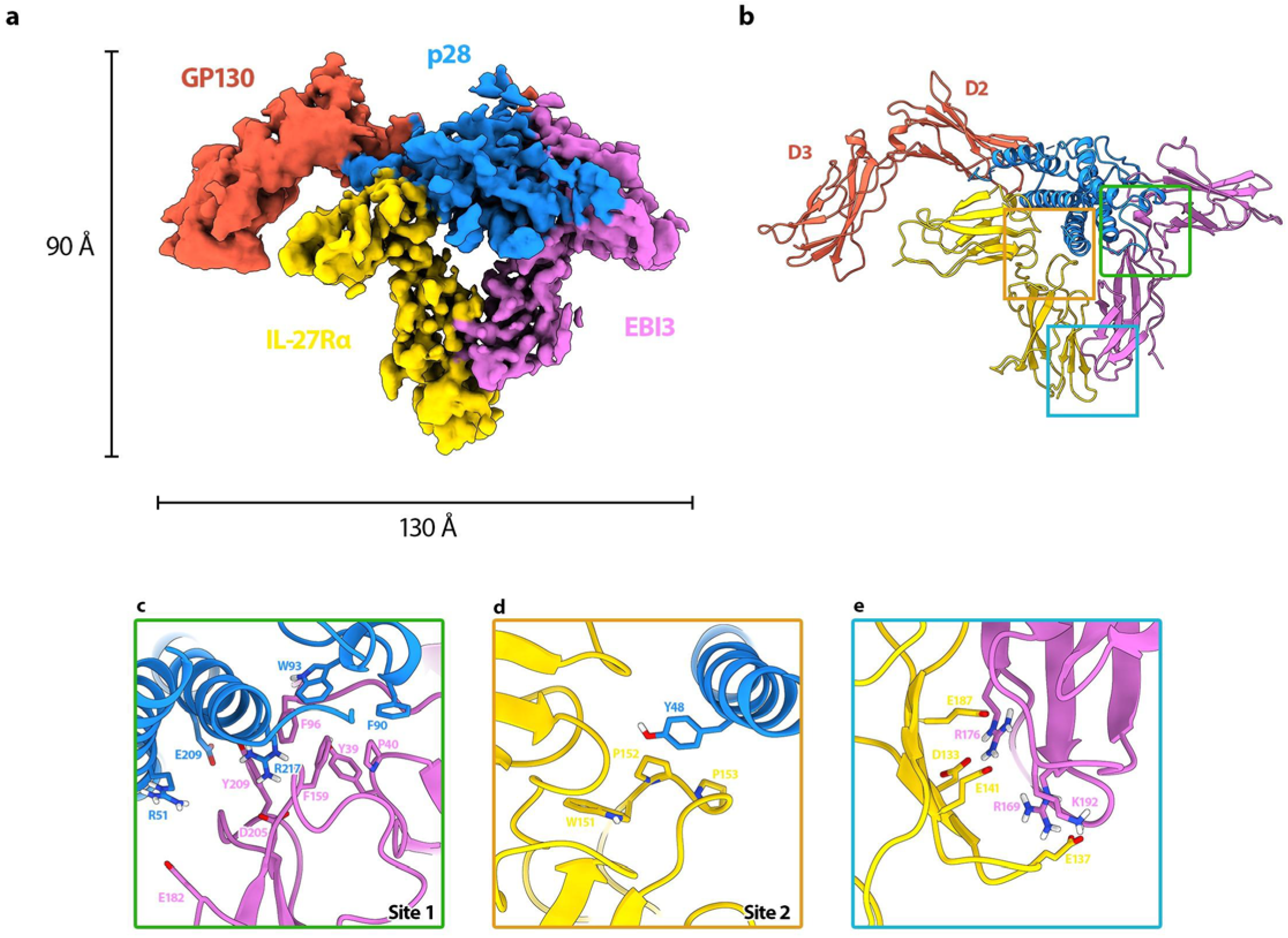
CryoEM reconstruction (**a**) and model (**b**) of the IL-27 receptor recognition complex. The complex consists of the IL-27 heterodimeric cytokine, p28 (blue) and EBI3 (purple), bound to two signalling receptors, GP130 (red) and IL-27Rα (yellow). EBI3 occupies site 1 on p28, while IL-27Rα occupies site 2. (**c-e**) Interaction interfaces of the complex. (**c**) The hinge between the two CHR domains of EBI3 form a hydrophobic groove composed of aromatic residues (Y39, P40, F96, F159 and Y209) that is filled by W93 of p28. Residue EBI3: D205, which is important for assembly of the heterodimeric cytokine ^28^, is also present in the binding interface and could form a salt bridge with p28:R217. (**d**) IL-27Rα binds site 2 of p28 at the apex of the elbow between its two CHR domains. The loops of IL-27Rα form an aromatic pocket comprised of residues IL-27Rα: W151, P152 and P153 in which the aromatic side chain of p28:Y48 slots into. (**e**) The orientation of IL-27Rα at site 2 is stabilized by a second interaction interface with EBI3, which is dominated by electrostatic complementarity between the two domains (IL-27Rα: E187, D133, E141, E137 and EBI3:R176, R169, K192). Key residues that likely mediate interactions at interfaces are shown as sticks.

The overall structure of the IL-27 cytokine recognition complex exhibits a classical architecture with a helical cytokine bundle central to the assembly (Fig. 1). For the IL-27 complex, the helical bundle is sandwiched by two ‘L’-shaped densities, analogous to other cytokine complexes. However, unlike previous structures, the IL-27 cytokine recognition complex has an additional prong that contacts the back face of the heterodimeric cytokine.

### Principle binding interfaces for IL27 cytokine recognition

Cytokines from the IL-12/IL-6 family have three highly-conserved principle binding interfaces: site 1, site 2 and site 3 ^23^. For the IL-27 recognition complex, site 1 is occupied by the second half of the heterodimeric cytokine, EBI3 (Figs. 1a-c). EBI3 is comprised of two Fibronectin type III (FNIII) domains bent into an ‘L’-shaped arrangement. Docking and minimal refinement of the EBI3 Alpha fold model into the map show that the bend of the EBI3 elbow is formed by a cluster of aromatic residues (EBI3: Y39, P40, F96, F159, and Y209), which create a hydrophobic groove recognized by p28:W93 (Fig. 1c). This feature of knobs and holes shape recognition is consistent with other cytokine binding interfaces ^23^. Our p28:EBI3 interface is supported by mutagenesis studies showing that equivalent residues in the human cytokine (EBI3:F97; p28:W97) are essential for cytokine complex formation ^28^. The same study identified an aspartic acid on EBI3 that also influences stability of the heterodimer ^28^. In our model, the equivalent murine residue (EBI3:D205) is nearby an arginine on p28 (p28:R217) and may form an additional salt bridge to stabilize the orientation of the heterodimer (Fig. 1c).

For the IL-6:GP130 cytokine recognition complex, site 2 of IL-6 is occupied by the two CHR domains of GP130 (D2 and D3) ^20^. To understand how GP130 recognizes chemically unique cytokines, we next investigated the IL-27 site 2 interface. By contrast to the IL-6 recognition complex, we find that site 2 of p28 is formed by the apex of the elbow region between the two CHR domains of IL-27Rα (Figs. 1a, b). Here we find that the knob and hole pattern is encoded by aromatic residues of IL-27Rα (IL-27Rα: P153, P152, and W151), which form a pocket to bind a tyrosine sidechain extending from p28 (p28:Y48) (Fig. 1d). In the IL-6:GP130 cytokine recognition complex, the equivalent tyrosine on IL-6 slots into a hydrophobic groove formed by the interface between D2 and D3 of GP130 ^20^.

The position of EBI3 at site 1 and IL-27Rα at site 2 are coordinated by a second interaction interface. EBI3 directly contacts the second CHR domain of IL-27Rα. By contrast to the hydrophobic knob and hole recognition motifs observed for p28, this secondary interface is dominated by electrostatic interactions (Figs. 1a, e, and Supplementary Fig. 4). Positively charged residues of EBI3 (EBI3: R169, R176, K192) face a negatively charged surface of IL-27Rα (IL-27Rα: D133, E137, E141, E187). These residues could be involved in a series of salt bridges that further contribute to the stability of the assembly.

The three N-terminal domains of GP130 are essential for cytokine recognition and signalling ^20,29^. A crystal structure of GP130 in complex with IL-6 showed that GP130 can engage cytokines through both site 2 and site 3 interactions ^20^. While the first two CHR domains of GP130 (D2 and D3) form the primary interface at site 2, the Ig domain (D1) of GP130 at site 3 bridges a second cytokine and is responsible for the lateral hexameric complex. In our structure of the IL-27 cytokine recognition complex, we observe a third prong of density extending from the core complex. Due to the poor angular distribution of particle orientations and resulting resolution anisotropy, we were unable to model this density. Based on our biochemical characterization of the complex showing that all components are present (Supplementary Fig.1), we conclude that some or all of this density could be accounted for by GP130. Indeed, biochemical data confirm that GP130 engages p28 at site 3^28^.

### Kinetic drivers of IL-27 signalling complex assembly

To understand the kinetic drivers underpinning assembly of the IL-27 signalling complex we defined the binding affinities for each of the subcomponents using surface plasmon resonance (SPR). First, we set out to identify which of the two signalling receptors bound the IL-27 heterodimer with higher affinity. We immobilized biotinylated IL-27Rα or GP130 on a streptavidin SPR surface and passed a range of IL-27 concentrations to measure rates of association and dissociation (Figs. 2 a,b). We found that IL-27Rα has a higher affinity (K _D_: 0.29 nM) for the heterodimeric cytokine than GP130 (K _D_: 3.1 nM), in agreement with biochemical data showing that GP130 engages the low-affinity binding site 3 ^28^. These results suggest a two-step binding mode for IL-27 complex formation, where IL-27 binds IL-27Rα with high affinity in a first step, and subsequently recruits GP130 with lower affinity to form the signalling complex. We next wanted to understand the roles of EBI3 and p28 in the kinetics of complex formation. As we were unable to produce recombinant EBI3, we focused our analysis on p28. We quantified the binding affinity of p28 for each signalling receptor immobilized to the streptavidin SPR chip (Fig. 2c, d). We observed only weak binding of p28 to IL-27Rα (K _D_: 2.8 μM) and no binding to GP130 at the doses tested, consistent with weak signalling responses elicited by p28 when compared to IL-27 in CD8 T cells (Fig. 2e, f). Taken together, these data support a stabilizing role of EBI3 in formation of the IL-27 signalling complex.

**Figure 2.**
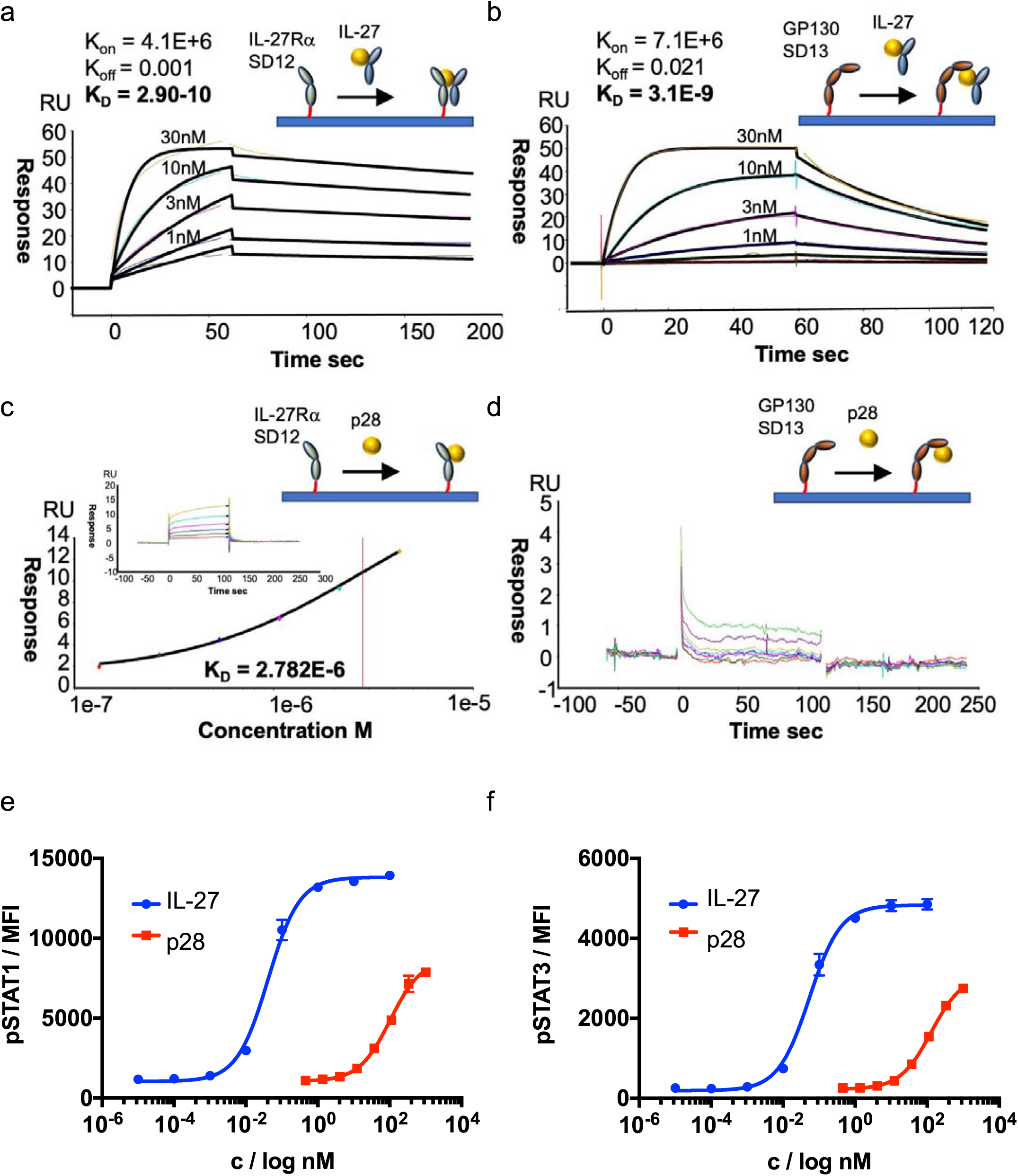
Biophysical analysis of the assembly of the IL-27 receptor complex. **(a-d)** For SPR measurements, IL-27Rα or GP130 were immobilized on the chip surface by biotin-streptavidin interaction and IL-27 or p28 were flowed across the chip in solution. (**a, b).** Kinetic charts for IL-27Rα (**a**) and GP130 (**b**) with inserts for affinity curves. Concentrations used are shown on the curves. K_on_, K_off_ and K_D_ values are shown. **(c, d).** Equilibrium charts for IL-27Rα (**c**) and GP130 (**d**) with inserts for affinity curves. Dose response curves for pSTAT1 **(e)** and pSTAT3 **(f)** in resting mouse CD8+ T cells. Cells were stimulated with mIL-27 or p28 for 15 minutes with the indicated doses. Data shown are the mean of four biological replicates with error bars depicting standard error of the mean.

## Discussion

IL-27 is a model system for signalling by heterodimeric cytokines, including IL-12 and IL-23. A comparison of the heterodimeric IL-23 receptor complex (PDB: 6WDQ) ^30^ with our structure shows that EBI3 and the IL-12β subunit (p40) overlay with site 1 (Fig. 3a). In addition, we observe that the GP130 density extending away from the core IL-27 complex has similarities to the IL-23 signalling receptor. The arrangement of these extracellular domains likely directs the orientation of intracellular regions that activate the JAK/STAT pathway. Therefore, it may be possible that the bend between D2 and D3 of GP130 in the signalling receptor is a topological requirement for activation by IL-12, IL-23 and IL-27.

**Figure 3.**
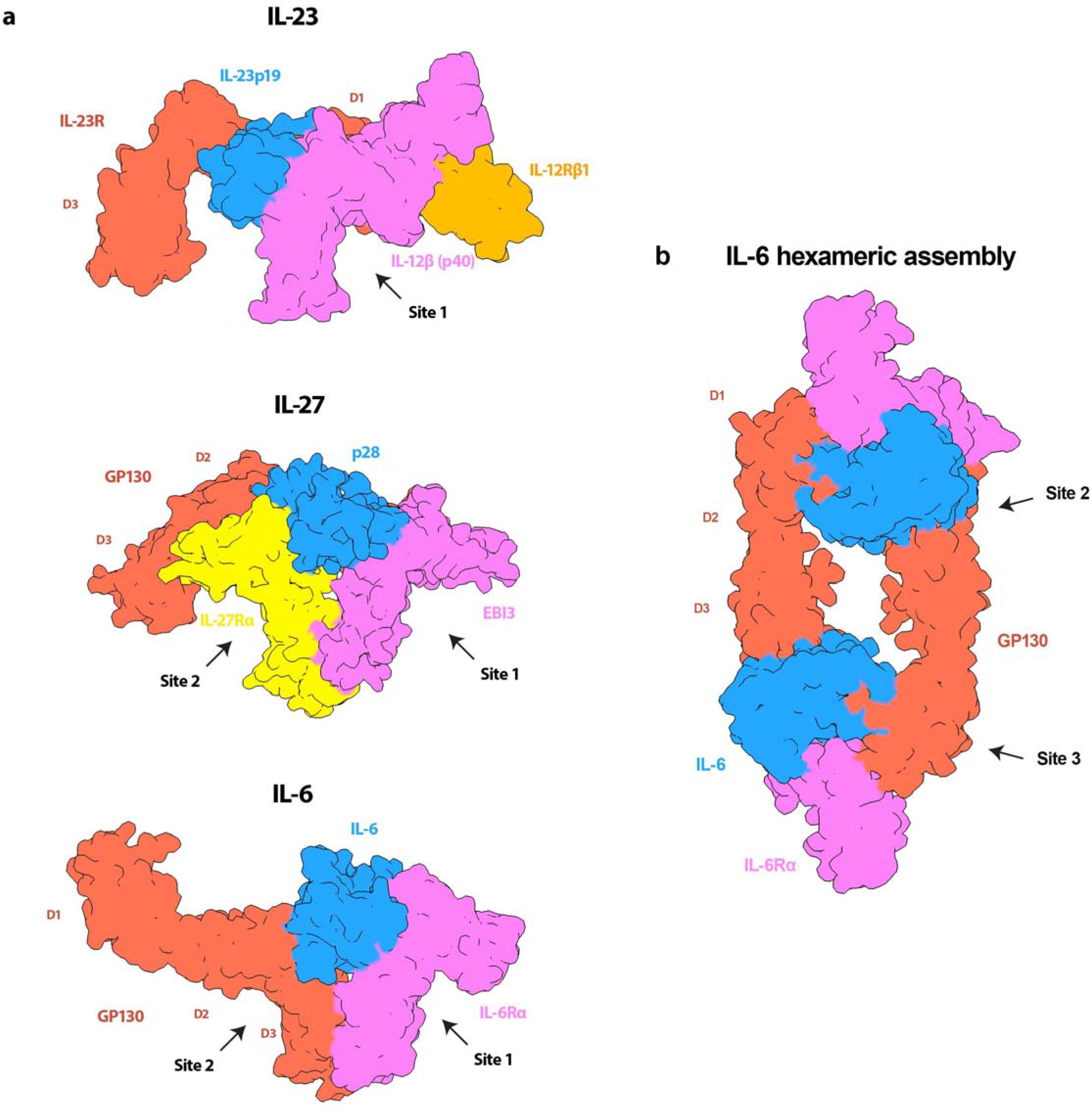
Model for how the IL-27 receptor recognition complex compares with IL-23 and IL-6 cytokine families. (**a**) (top panel) Surface rendering of the IL-23 receptor recognition complex (PDB: 6WDQ) coloured according to protein components: IL-23 receptor (red), IL-12β (p40) (purple), IL-23p19 (blue), IL-12Rβ1 (orange). (middle panel) A comparison with IL-27 receptor complex shows that both GP130 (red) and IL-23 receptor protrude from the core cytokines, orienting domains similarly towards the membrane. Both EBI3 (purple) and the other component of the IL-23 heterocytokine, p40, occupy the site 1 epitope. (bottom panel) Surface rendering of the IL-6 receptor recognition complex (PDB: 1P9M) colored according to protein components: IL-6 (blue), IL-6Rα (purple), GP130 (red). (middle panel) A comparison with the IL-27 receptor complex shows that the nonsignaling components of both complexes (EBI3 and IL-6Rα) occupy site 1, while GP130 binds in a different way. For IL-6, GP130 binds site 2 through an interaction site located between the CHR domains, D2 and D3. While site 2 is of IL-27 is occupied by IL-27Rα. **(b)** Surface rendering of the hexameric IL-6 assembly (PDB: 1P9M) colored according to protein components: IL-6 (blue), IL-6Rα (purple), GP130 (red). GP130 bound to site 2 of one IL-6 molecule bridges a second cytokine by binding at site 3 to stabilize the complex.

IL-27 is also a central member of the IL-6 family of cytokines that signal through GP130 ^31^. The diversity in cellular responses within the family stems in part from the strict transcriptional control of cytokines secreted by different cell types ^32^. Although IL-27 is a heterodimeric cytokine, there are some instances whereby the two components are independently expressed ^33,34^. In the absence of EBI3, p28 can act as an antagonist for IL-6-mediated GP130 signalling ^13^. Our biophysical data show that binding of p28 alone to GP130 is negligible (Fig. 2). A comparison of the IL-6 cytokine recognition complex (PDB:1P9M) with our structure shows that binding of EBI3 to site 1 overlaps with site 1 of the IL-6 (Fig. 3a). Previous studies have reported that p28 can bind IL-6Rα ^17^. Though signalling by the putative p28:IL-6Rα is weaker compared to IL-6 or IL-27, it may be possible that IL-6Rα engages p28 at site 1 with low affinity to produce weak agonistic or antagonistic activities in certain contexts.

Chain sharing is a common theme among cytokine receptors ^9^ and contributes to the layers of complexity derived from relatively few building blocks. Although both IL-27 and IL-6 engage GP130, they do so through different interaction interfaces. One important difference between these two structures occurs at site 2 (Fig. 3a). While the two CHR domains of GP130 (D2 and D3) bind IL-6 at site 2, the equivalent position in the IL-27 system is occupied by IL-27Rα. Unlike the IL-6 co-receptor (IL-6Rα), the intracellular domains of GP130 and IL-27Rα both associate with JAKs proteins and activate signalling upon dimer formation. In the case of IL-27, the cognate co-receptor IL-27Rα could fill an analogous role to GP130 at site 2 in the IL-6-signaling complex. In both systems site 1 is occupied by non-signalling proteins: EBI3 and IL-6Rα. Our structural and biophysical data support role for auxiliary proteins at site 1 to enhance the stability of the signalling complex, while receptors occupying site 2 are responsible for signal transduction.

Signalling through the JAK/STAT pathway requires dimerization of signalling receptors. How that dimerization is achieved, and how the geometry of the signalling complex contributes to the functional diversity of the cascade, remains an open question. There is a wide range of oligomeric assemblies observed for GP130-cytokine complexes ^20,23^. For IL-6, dimerization of GP130 occurs through a hexameric assembly (2 copies of the IL-6:IL-6Rα:GP130 complex). Within the hexamer, GP130 bound to IL-6 site 2 dimerizes with a second copy bound to site 3 of the additional cytokine (Fig. 3b). In the IL-27 signalling complex, dimerization of IL-27Rα and GP130 may occur from receptors occupying sites 2 and 3 from the same cytokine, though additional cytokine-receptor stoichiometries cannot be ruled out. Indeed, the primary interaction interface observed in the IL-6:GP130 complex occurs at the apex of a bend between GP130 CHR domains (D2 and D3). In the IL-27 complex, this region is likely unoccupied and available to engage other cytokines. In addition, IL-27 has been shown to inhibit signalling by oncostatin M (OSM), an IL-6 family cytokine ^35^. Based on our structure, IL-27 immunosuppressant activities may be achieved by blocking inflammatory cytokines such as OSM from binding GP130 at site 2 in a signalling competent conformation. Understanding how mixtures of cytokines influence signalling assemblies and downstream cascades will inform new strategies in protein engineering for fine-tune cytokine responses.

## Material and methods

### Protein expression and purification

Murine IL-27 was cloned as a linker-connected single-chain variant (p28+EBI3) as described in ^36^ (Supplementary Fig. 1a). In the text we refer to this variant only as IL-27. Murine IL-27Rα (amino acids 28-224) and GP130 (amino acid 23-319) ectodomains were cloned and expressed as described in ^37^. Briefly, protein sequences were cloned into the pAcGP67-A vector (BD Biosciences) in frame with an N-terminal gp67 signal sequence, driving protein secretion, and a C-terminal hexahistidine tag. Baculovirus stocks were produced by transfection and amplification in *Spodoptera frugiperda* (*Sf*9) cells grown in SF900III media (Invitrogen) and protein expression was carried out in suspension *Trichoplusiani ni* (High Five) cells grown in InsectXpress media (Lonza).

Protein purification was carried out using the method described in ^38^. Hi-Five cells were pelleted with centrifugation at 2000 rpm, and impurities from the remaining media were removed by a precipitation step through addition of Tris pH 8.0, CaCl_2_ and NiCl_2_ to final concentrations of 200mM, 50mM and 1mM respectively. The precipitate formed was then removed through centrifugation at 6000 rpm. Nickel-NTA agarose beads (Qiagen) were added to the clarified media and the target proteins purified through batch binding followed by column washing in HBS buffer for murine IL-27Rα and GP130 or HBS-Hi buffer (HBS buffer supplemented to 500mM NaCl and 5% glycerol, pH 7.2) for murine IL-27sc. Elution was performed using HBS or HBS-Hi buffer plus 200mM imidazole. Final purification was performed by size exclusion chromatography on an ENrich SEC 650 10 x 300 column (Biorad), equilibrated in HBS or HBS-Hi buffers. Concentration of the purified sample was carried out using 30kDa Millipore Amicon-Ultra spin concentrators. Recombinant proteins were purified to greater than 98% homogeneity. For cryoEM studies, IL-27Rα, GP130 and IL-27 were mixed in a 1:1:1 ratio and subsequently purified by size-exclusion chromatography.

To generate biotinylated proteins for surface plasmon resonance studies the GP130 sequence was subcloned into the pAcGP67-A vector with a C-terminal biotin acceptor peptide (BAP)–LNDIFEAQKIEWHW followed by a hexa-histidine tag. The purified protein was biotinylated with BirA ligase following a previously described protocol ^38^. IL-27Rα was N-terminal biotinylated in vitro using EZ-Link Sulfo-NHS-SS-Biotin (Pierce) at pH 6.5.

### Surface plasmon resonance

Surface plasmon resonance was used to determine the binding affinity of the recombinantly produced IL-27 and p28 to IL-27Rα and GP130. Biotinylated IL-27Rα and GP130 were immobilised onto the chip surface with streptavidin. Series S Sensor SA (GE Healthcare) chips were primed in 10 mM HEPES, 150 mM NaCl, 0.02% TWEEN-20, prior to immobilisation of the biotinylated receptor. Analysis runs were performed in 10 mM HEPES, 150 mM NaCl, 0.05% TWEEN-20 and 0.5% BSA. A Biacore T100 (T200 Sensitivity Enhanced) was used for measurement and Biacore T200 Evaluation Software 3.0 was used for data analysis.

### Phospho-flow analysis

For phospho-flow analysis of STAT1 and STAT3, mouse CD8 T cells purify from a wildtype spleen were plated at a density of 2 x 10^5^ cells per well in 50 μL in a 96-well V bottom plate. Cells were left unstimulated or simulated with 3-fold serially diluted IL-27sc or p28 variants (50 μL per well) for 15 minutes at 37°C before fixation with 2% paraformaldehyde for 10 minutes at room temperature. Cells were washed in PBS and permeabilised in ice-cold 100% methanol and incubated on ice for a minimum of 30 minutes. Cells were fluorescently barcoded as previously described ^37,39^. Briefly, a panel of 16 combinations of two NHS-dyes (Pacific Blue and DyLight800, Thermo) was used to stain individual wells on ice for 35 minutes before stopping the reaction by washing in PBS/0.5% BSA. Once barcoded, the 16 populations were pooled together for antibody staining. Cells were stained with anti-pSTAT3^Alexa488^ (Biolegend #651006) and anti-pSTAT1^Alexa647^ (Cell Signalling Technologies #8009). During acquisition, individual populations were identified according to the barcoding pattern and pSTAT3^Alexa488^ and pSTAT1^Alexa647^ MFI was quantified for all populations. MFI was plotted and sigmoidal dose response curves were fitted using Prism software (Version 7, GraphPad).

### CryoEM sample preparation and data collection

Pre-formed IL-27:IL-27Rα:GP130 complex sample was diluted to 0.1 mg/ml and vitrified in liquid ethane using Vitrobot Mark IV (Thermo Fisher Scientific). Lacey Carbon Au 300 grids (EMS) were glow discharged in air using a Cressington 208 for 60 seconds prior to application of the sample. 4μL of sample were applied to the grids at a temperature of 20°C and a humidity level of 100%. Grids were then immediately blotted (force −2, time 2 seconds) and plunge-frozen into liquid ethane cooled to liquid nitrogen temperature.

Grids were imaged using a 300kV a Titan Krios transmission electron microscope (Thermo Fisher Scientific) equipped with K3 camera (Gatan) operated in super resolution mode. Movies were collected at 81,000x magnification and binned by two on the camera (calibrated pixel size of 1.06 Å/pixel). Images were taken over a defocus range of −0.5 μm to −2.25 μm with a total accumulated dose of 50 electrons per Å^2^ using EPU (Thermo Fisher Scientific, version 2.11.1.11) automated data software. A summary of imaging conditions is presented in Supplementary Table 1.

### CryoEM data processing

28,437 micrographs were pre-processing using cryoSPARC (v.3.3.1) ^26^ patch motion correction and patch CTF. The datasets were manually curated to remove movies with substantial drift and crystalline ice, resulting in a total of 17,228 micrographs going on for further processing. Deep learning models used in Topaz (v.0.2.5) ^40^ were used to automatically pick particles from these micrographs within the cryoSPARC workflow. We used the Topaz pre-trained model (ResnNet16) for particle picking in the first instance. During the initial 2D classification, rare views were combined and used as input for a second round of Topaz training. Particles from both rounds were combined and duplicates were removed. A total of 3,575,367 particles were extracted at 2.12 Å/pixel (bin by 2) and subject to iterative 2D classification to remove ice contamination, carbon edges and broken particles. Of these, 628,386 particles were used to create an initial model using cryoSPARC’s ab initio reconstruction procedure with three classes. By comparing the reconstructions, it was obvious that one class represented a subcomponent of the complex. Particles from the other two classes (486,648 particles) showing similar structural features were merged and refined using nonuniform refinement. Particles were then re-extracted at 1.06 Å/pixel and movement within the ice was corrected using cryoSPARCs own implementation of local motion correction. The dataset was then subjected to heterogenous refinement using six classes. All six classes were individually refined using homogeneous refinement. The three classes that refined to the highest resolution were pooled (289,489 particles) and were further refined using local refinement procedures manually centered on the core. The final map resolution 3.8 Å was assessed using the gold standard FSC at a threshold of 0.143 and locally filtered using cryoSPARCs own implementation.

### Model building and refinement

An initial model of IL-27 was generated by rigid body fitting each protein into the locally sharpened map generated by DeepEMhancer ^41^. Rigid body fitting was performed in Chimera ^42^. The models used were murine p28 (AlphaFold entry ID: Q8K3I6), murine IL-27Rα (AlphaFold entry ID: O703940), murine EBI3 (AlphaFold entry ID: O35228), and murine GP130 (AlphaFold entry ID: Q00560). Alpha fold models were trimmed to reflect the domain boundaries in the constructs used.

The initial model of IL-27 was refined into the cryoEM map using ISOLDE (v.1.3) ^43^ implemented in ChimeraX (v.1.3) ^44^. During refinement we applied adaptive distance restraints to each subunit. A loop region of p28 (residues 178-194) was removed to prevent steric clashes with GP130. The atomics models were refined using phenix.real_space_refine in Phenix (v.1.20.1-4487) ^45^ with secondary structure, reference model, and geometry restraints. B-factors were refined in Phenix. Model FSC validation tools and the overall quality of the model were assessed in the map using the cryoEM validation tools in Phenix and MolProbity ^46^ (Supplementary Table 1).

## Data Availability

Data supporting the findings of this manuscript are available from the corresponding authors upon reasonable request. The accession numbers for the EM map and models of the IL27-receptor recognition complex reported in this paper are XXX and XXX.

## Acknowledgments

We thank members of the Moraga and Bubeck laboratories for helpful advice and discussion. This work was supported by the Wellcome-Trust-202323/Z/16/Z (I.M.), ERC-206-STG grant (I.M. P.K.F., S.W.). We thank S. Islam for computational support and P. Simpson for EM support. Initial screening of samples was carried out at Imperial College London Centre for Structural Biology; cryoEM data was collected at Diamond Light Source. We thank Diamond for access and support of the Cryo-EM facilities at the UK national electron bio-imaging centre (eBIC), proposal BI25127, funded by the Wellcome Trust, MRC and BBSRC. We also wish to thank R. Sundaramoorthy for his assistance in the initial stages of the project.

## Author Contributions

Y.J. conducted cryoEM work. S.G. and D.B. built and refined atomic models of the complex. P.K.F. recombinantly expressed proteins and perform SPR studies. S.W. conducted signalling studies. D.B. and I.M. conceived the ideas. Y.J., S.G and D.B. analyzed cryoEM data. D.B. and I.M. wrote the manuscript. S.G. and Y.J. generated the figures. All authors assisted with manuscript editing.

## Competing Interests Statement

The authors declare that there are no competing interests.

## Notes

### Competing Interest Statement

The authors have declared no competing interest.

## References

1. Stumhofer, J.S. et al. Interleukin 27 negatively regulates the development of interleukin 17-producing T helper cells during chronic inflammation of the central nervous system. Nat Immunol 7, 937–45 (2006).

2. Diveu, C. et al. IL-27 blocks RORc expression to inhibit lineage commitment of Th17 cells. J Immunol 182, 5748–56 (2009).

3. Yang, J. et al. Epstein-Barr virus-induced gene 3 negatively regulates IL-17, IL-22 and RORgamma t. Eur J Immunol 38, 1204–14 (2008).

4. Hall, A.O. et al. The cytokines interleukin 27 and interferon-gamma promote distinct Treg cell populations required to limit infection-induced pathology. Immunity 37, 511–23 (2012).

5. Stumhofer, J.S. et al. Interleukins 27 and 6 induce STAT3-mediated T cell production of interleukin 10. Nat Immunol 8, 1363–71 (2007).

6. Chihara, N. et al. Induction and transcriptional regulation of the co-inhibitory gene module in T cells. Nature 558, 454–459 (2018).

7. DeLong, J.H. et al. IL-27 and TCR Stimulation Promote T Cell Expression of Multiple Inhibitory Receptors. Immunohorizons 3, 13–25 (2019).

8. Jia, H., Dilger, P., Bird, C. & Wadhwa, M. IL-27 Promotes Proliferation of Human Leukemic Cell Lines Through the MAPK/ERK Signaling Pathway and Suppresses Sensitivity to Chemotherapeutic Drugs. J Interferon Cytokine Res 36, 302–16 (2016).

9. Vignali, D.A. & Kuchroo, V.K. IL-12 family cytokines: immunological playmakers. Nat Immunol 13, 722–8 (2012).

10. Kourko, O., Seaver, K., Odoardi, N., Basta, S. & Gee, K. IL-27, IL-30, and IL-35: A Cytokine Triumvirate in Cancer. Front Oncol 9, 969 (2019).

11. Tait Wojno, E.D., Hunter, C.A. & Stumhofer, J.S. The Immunobiology of the Interleukin-12 Family: Room for Discovery. Immunity 50, 851–870 (2019).

12. Trinchieri, G., Pflanz, S. & Kastelein, R. The IL-12 family of heterodimeric cytokines: New players in the regulation of T-cell responses. Immunity 19, 641–644 (2003).

13. Stumhofer, J.S. et al. A role for IL-27p28 as an antagonist of gp130-mediated signaling. Nat Immunol 11, 1119–26.

14. Collison, L.W. et al. The composition and signaling of the IL-35 receptor are unconventional. Nat Immunol 13, 290–9.

15. Crabe, S. et al. The IL-27 p28 subunit binds cytokine-like factor 1 to form a cytokine regulating NK and T cell activities requiring IL-6R for signaling. J Immunol 183, 7692–702 (2009).

16. Wang, R.X., Yu, C.R., Mahdi, R.M. & Egwuagu, C.E. Novel IL27p28/IL12p40 cytokine suppressed experimental autoimmune uveitis by inhibiting autoreactive Th1/Th17 cells and promoting expansion of regulatory T cells. J Biol Chem 287, 36012–21 (2012).

17. Garbers, C. et al. An interleukin-6 receptor-dependent molecular switch mediates signal transduction of the IL-27 cytokine subunit p28 (IL-30) via a gp130 protein receptor homodimer. J Biol Chem 288, 4346–54 (2013).

18. Grotzinger, J., Kurapkat, G., Wollmer, A., Kalai, M. & Rose-John, S. The family of the IL-6-type cytokines: specificity and promiscuity of the receptor complexes. Proteins 27, 96–109 (1997).

19. Hunter, C.A. & Jones, S.A. IL-6 as a keystone cytokine in health and disease. Nat Immunol 16, 448–57 (2015).

20. Boulanger, M.J., Chow, D.C., Brevnova, E.E. & Garcia, K.C. Hexameric structure and assembly of the interleukin-6/IL-6 alpha-receptor/gp130 complex. Science 300, 2101–4 (2003).

21. Boulanger, M.J., Bankovich, A.J., Kortemme, T., Baker, D. & Garcia, K.C. Convergent mechanisms for recognition of divergent cytokines by the shared signaling receptor gp130. Mol Cell 12, 577–89 (2003).

22. Skiniotis, G., Lupardus, P.J., Martick, M., Walz, T. & Garcia, K.C. Structural organization of a full-length gp130/LIF-R cytokine receptor transmembrane complex. Mol Cell 31, 737–48 (2008).

23. Wang, X., Lupardus, P., Laporte, S.L. & Garcia, K.C. Structural biology of shared cytokine receptors. Annu Rev Immunol 27, 29–60 (2009).

24. Huyton, T. et al. An unusual cytokine:Ig-domain interaction revealed in the crystal structure of leukemia inhibitory factor (LIF) in complex with the LIF receptor. Proc Natl Acad Sci U S A 104, 12737–42 (2007).

25. Wilmes, S. et al. Competitive binding of STATs to receptor phospho-Tyr motifs accounts for altered cytokine responses. Elife 10(2021).

26. Punjani, A., Rubinstein, J.L., Fleet, D.J. & Brubaker, M.A. cryoSPARC: algorithms for rapid unsupervised cryo-EM structure determination. Nat Methods 14, 290–296 (2017).

27. Jumper, J. et al. Highly accurate protein structure prediction with AlphaFold. Nature 596, 583–589 (2021).

28. Rousseau, F. et al. IL-27 structural analysis demonstrates similarities with ciliary neurotrophic factor (CNTF) and leads to the identification of antagonistic variants. Proc Natl Acad Sci U S A 107, 19420–5 (2010).

29. Dahmen, H. et al. Activation of the signal transducer gp130 by interleukin-11 and interleukin-6 is mediated by similar molecular interactions. Biochem J 331 (Pt 3), 695–702 (1998).

30. Glassman, C.R. et al. Structural basis for IL-12 and IL-23 receptor sharing reveals a gateway for shaping actions on T versus NK cells. Cell 184, 983–999 e24 (2021).

31. Hunter, C.A. & Kastelein, R. Interleukin-27: balancing protective and pathological immunity. Immunity 37, 960–9 (2012).

32. Taga, T. & Kishimoto, T. Gp130 and the interleukin-6 family of cytokines. Annu Rev Immunol 15, 797–819 (1997).

33. Devergne, O. et al. A novel interleukin-12 p40-related protein induced by latent Epstein-Barr virus infection in B lymphocytes. J Virol 70, 1143–53 (1996).

34. Maaser, C., Egan, L.J., Birkenbach, M.P., Eckmann, L. & Kagnoff, M.F. Expression of Epstein-Barr virus-induced gene 3 and other interleukin-12-related molecules by human intestinal epithelium. Immunology 112, 437–45 (2004).

35. Baker, B.J., Park, K.W., Qin, H., Ma, X. & Benveniste, E.N. IL-27 inhibits OSM-mediated TNF-alpha and iNOS gene expression in microglia. Glia 58, 1082–93 (2010).

36. Oniki, S. et al. Interleukin-23 and interleukin-27 exert quite different antitumor and vaccine effects on poorly immunogenic melanoma. Cancer Res 66, 6395–404 (2006).

37. Martinez-Fabregas, J. et al. Kinetics of cytokine receptor trafficking determine signaling and functional selectivity. Elife 8(2019).

38. Spangler, J.B., Moraga, I., Jude, K.M., Savvides, C.S. & Garcia, K.C. A strategy for the selection of monovalent antibodies that span protein dimer interfaces. J Biol Chem 294, 13876–13886 (2019).

39. Krutzik, P.O. & Nolan, G.P. Fluorescent cell barcoding in flow cytometry allows high-throughput drug screening and signaling profiling. Nat Methods 3, 361–8 (2006).

40. Bepler, T., Kelley, K., Noble, A.J. & Berger, B. Topaz-Denoise: general deep denoising models for cryoEM and cryoET. Nat Commun 11, 5208 (2020).

41. Sanchez-Garcia, R. et al. DeepEMhancer: a deep learning solution for cryo-EM volume post-processing. Commun Biol 4, 874 (2021).

42. Pettersen, E.F. et al. UCSF Chimera--a visualization system for exploratory research and analysis. J Comput Chem 25, 1605–12 (2004).

43. Croll, T.I. ISOLDE: a physically realistic environment for model building into low-resolution electron-density maps. Acta Crystallogr D Struct Biol 74, 519–530 (2018).

44. Pettersen, E.F. et al. UCSF ChimeraX: Structure visualization for researchers, educators, and developers. Protein Sci 30, 70–82 (2021).

45. Afonine, P.V. et al. New tools for the analysis and validation of cryo-EM maps and atomic models. Acta Crystallogr D Struct Biol 74, 814–840 (2018).

46. Williams, C.J. et al. MolProbity: More and better reference data for improved all-atom structure validation. Protein Sci 27, 293–315 (2018).

