## Supplemental Figures and Tables for "Structural insights into the assembly and activation of IL-27 signalling complex" (from another version of this preprint: version 1, 2022-02-19)

**ONLINE SUPPLEMENTARY INFORMATION**

**SUPPLEMENTARY TABLES**

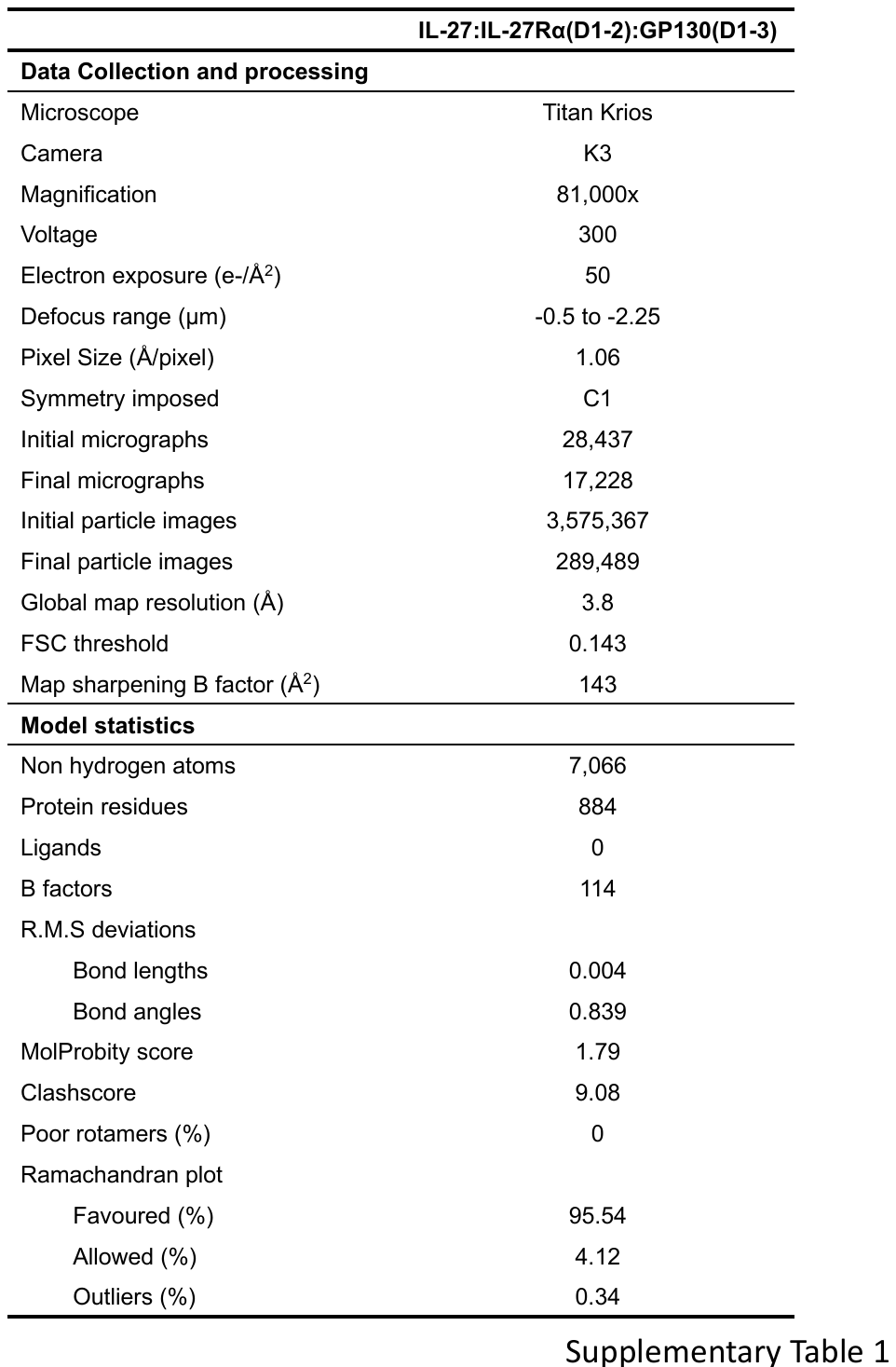

**Supplementary Table 1. CryoEM data collection and model validation statistics**

**SUPPLEMENTARY FIGURES**

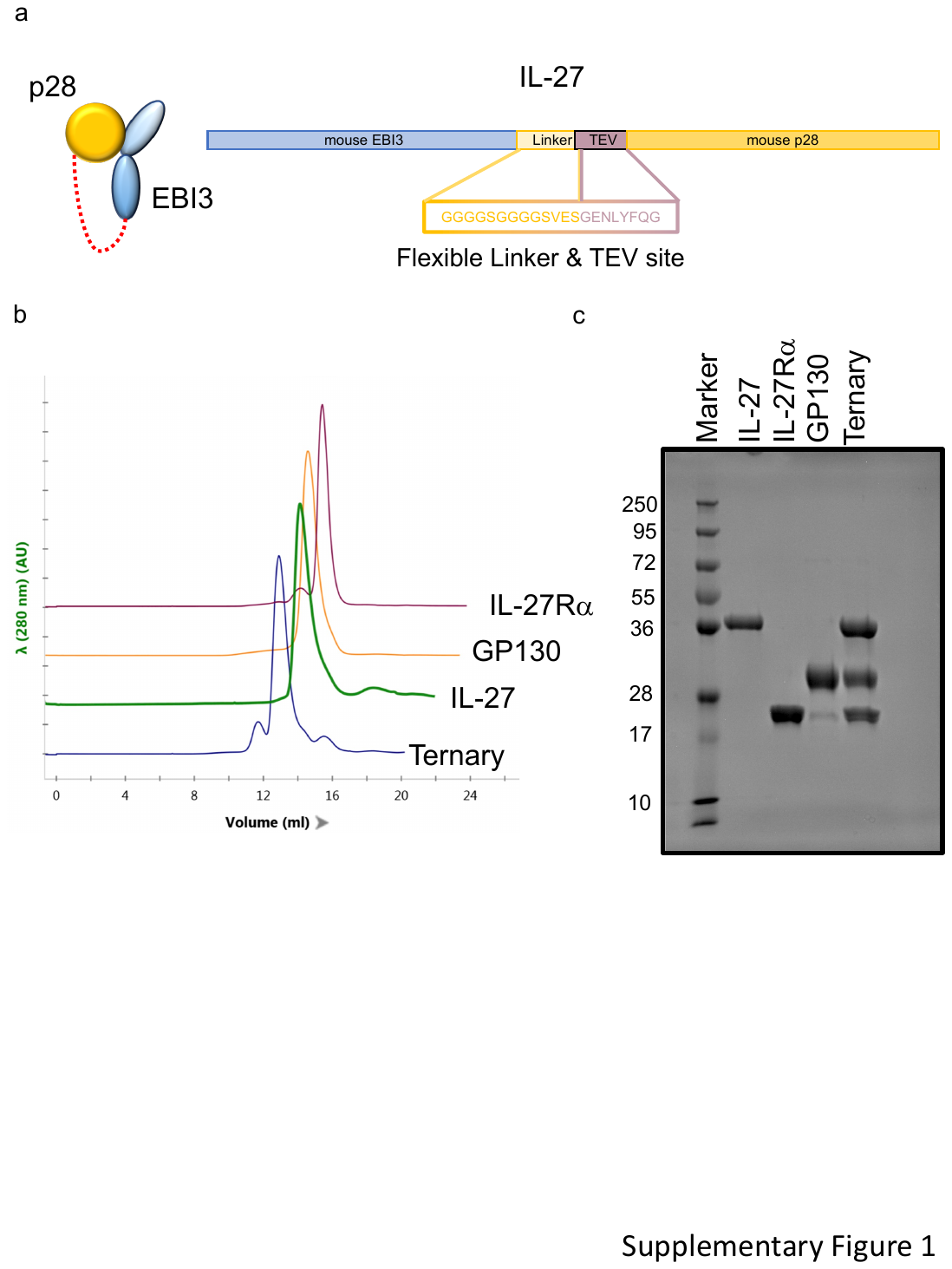

**Supplementary Figure 1.**  **Assembly and characterization of the IL-27-receptor complex.** **(a)** Schematic representation of the single chain IL-27 heterodimer. **(b)** Chromatogram overlay from size exclusion chromatography of the different IL-27 complex components. **(c)** Coomassie-stained SDS-PAGE analysis under reducing conditions of the individual components plus the IL-27:IL-27Rα:GP130 complex used for cryoEM studies.

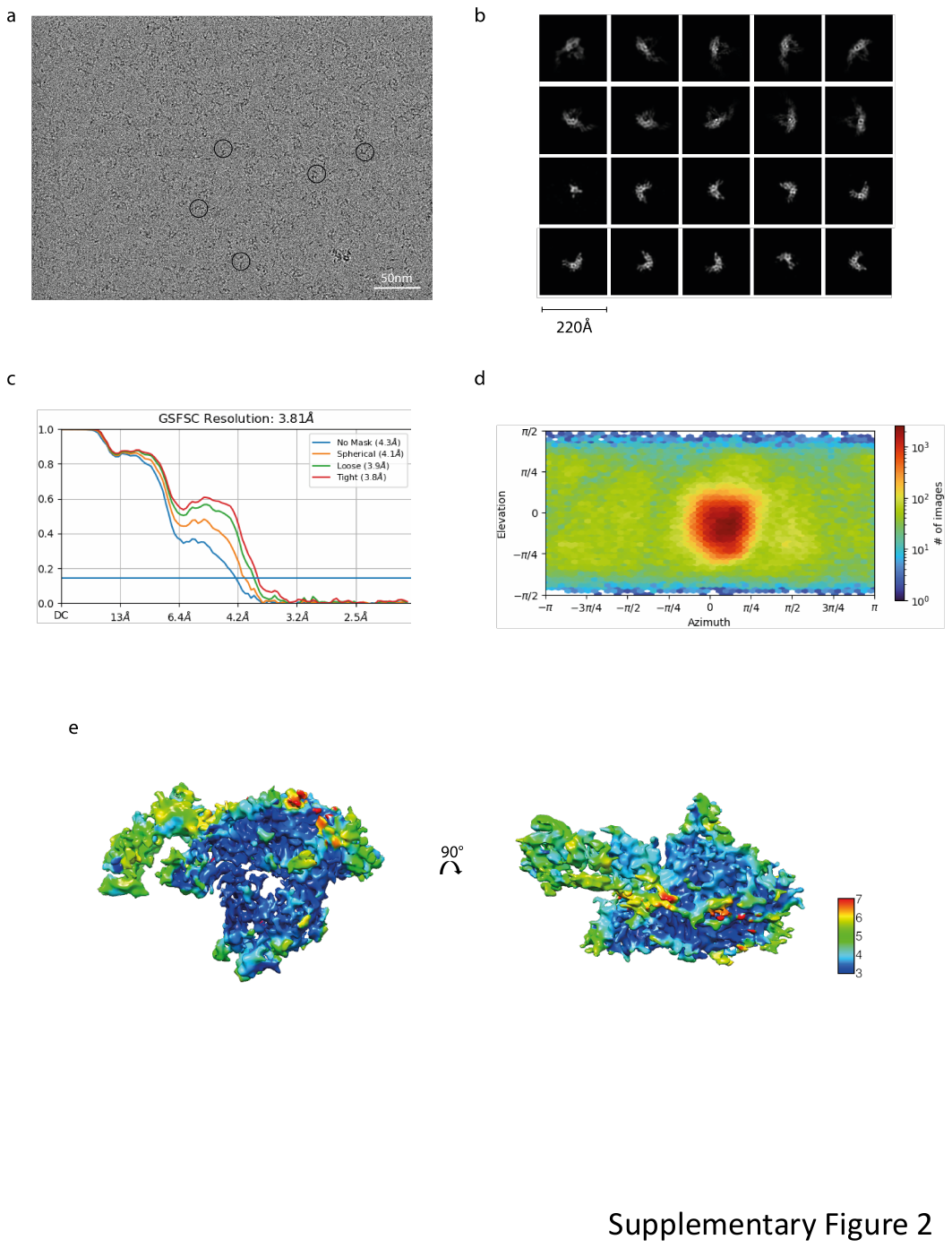

**Supplementary Figure 2.** Supplementary cryoEM data. **(a)** Raw cryoEM micrograph with examples of the IL-27 receptor complex circled. Scale bar 50 nm **(b)** 2D class averages. Scale bar 220Å. **(c)** Gold standard Fourier shell correlation (GSFSC) for the final reconstruction. The resolution at the 0.143 cutoff is reported. **(d)** Angular distribution plot. **(e)** IL-27 receptor recognition reconstruction filtered according to local resolution ranging from 3 Å (blue) to 7 Å (red).

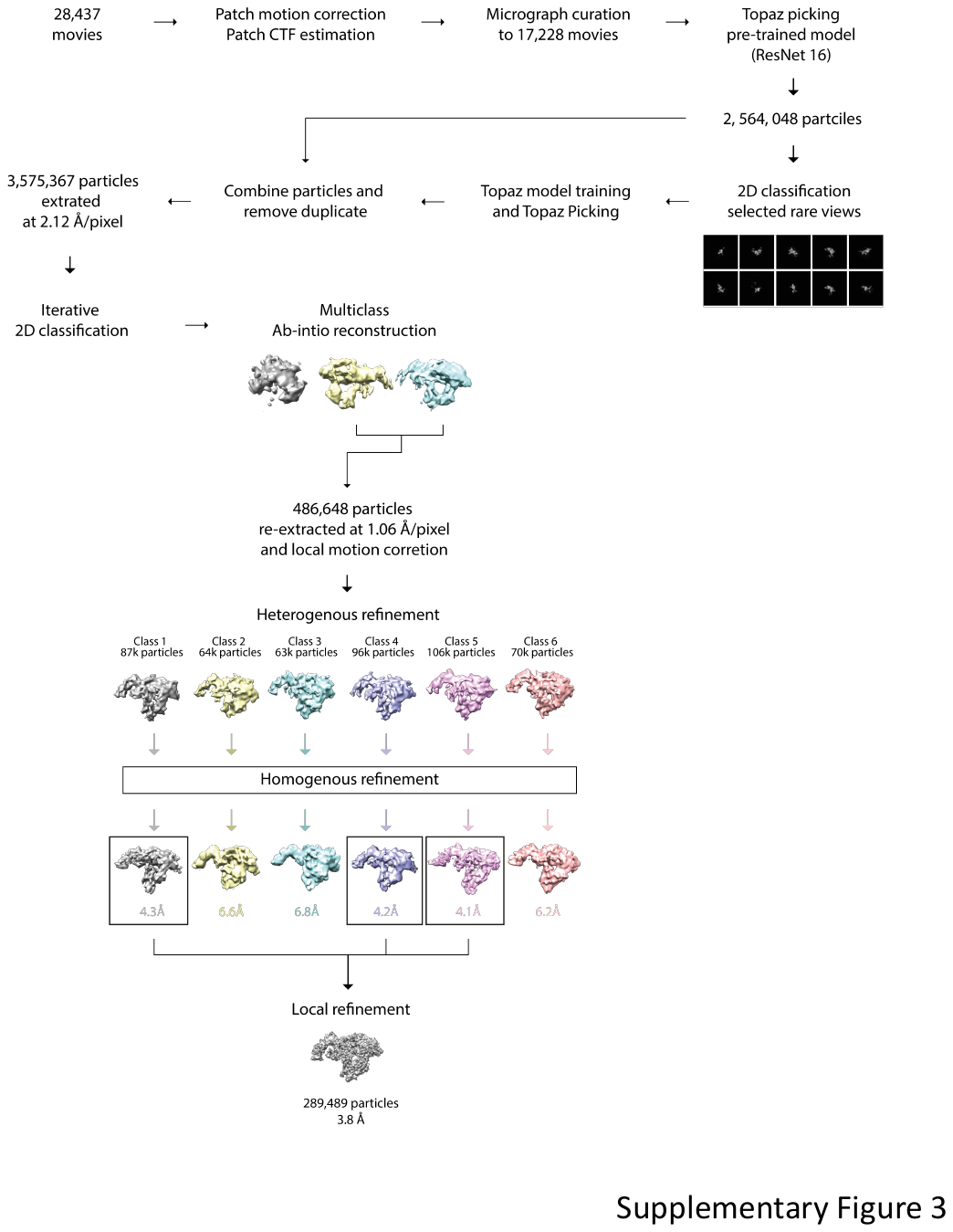

**Supplementary Figure 3.** CryoEM image processing workflow outlining steps performed to obtain the structure of IL-27 heterodimer complex. All processing was performed using CryoSPARC v.3.3.1 (see Methods for details).

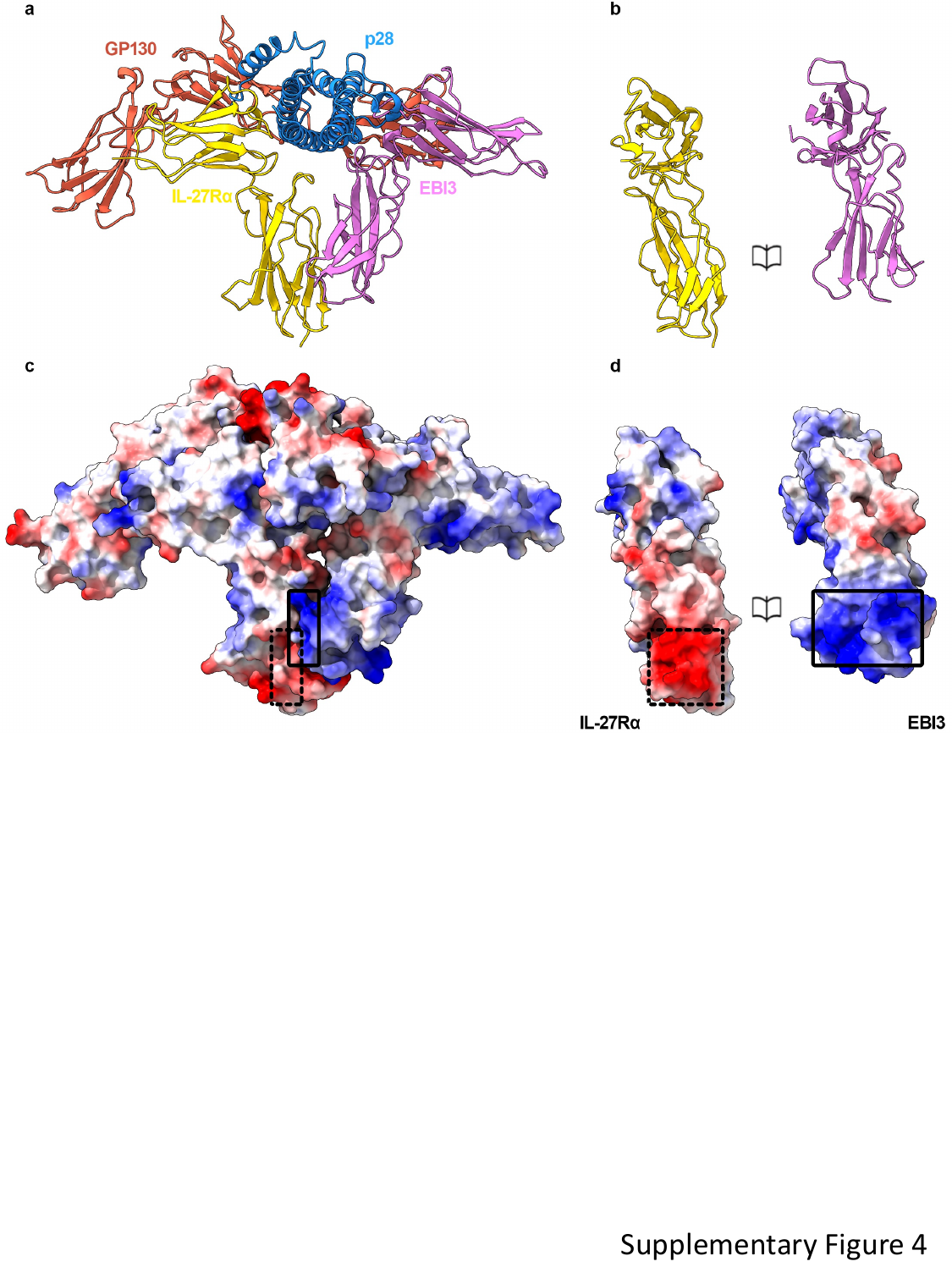

**Supplementary Figure 4.** EBI3:IL27-Rα interface. **(a)** Ribbon representation of the IL-27 receptor recognition complex. Individual proteins colored: EBI3 (purple), IL-27Rα (yellow), p28 (blue), GP130 (red). **(b)** Book representation of the EBI3:IL-27Rα interface. **(c)** Coulombic electrostatic potential ranging from -10 (red) to 10 (blue) kcal/(mol·e) calculated from the models in **a**. **(d)** Book representation of the interface highlighting the contact area on EBI3 (black square) and IL-27Rα (dotted lines).
